# PDBe-SIFTS: an open-source tool for Structure Integration with Function, Taxonomy, and Sequences, featuring improved alignment, scoring scheme, and accelerated search

**DOI:** 10.64898/2026.04.30.721839

**Authors:** Adam Bellaiche, Preeti Choudhary, Sreenath Nair, Deborah Harrus, Conny Wing-Heng Yu, Syed Ahsan Tanweer, Genevieve Laura Evans, Stephanie W Lo, Maria J Martin, Jennifer R Fleming, Sameer Velankar

## Abstract

Structure Integration with Function, Taxonomy and Sequences (SIFTS) provides residue-level mappings between UniProt Knowledgebase sequences and Protein Data Bank structures and has historically been generated through internal Protein Data Bank in Europe (PDBe) pipelines. Here, PDBe-SIFTS is presented as a fully open-source, locally deployable implementation of this mapping framework. The pipeline combines fast, scalable sequence search using MMseqs2, an improved bounded scoring scheme for ranking candidate mappings, and residue-level mapping refinement based on backbone connectivity. PDBe-SIFTS is distributed as a Python package with command-line tools for 1) building a sequence search database, 2) identifying the best sequence-structure match, 3) one-to-one mapping at the residue level, and 4) generating SIFTS annotations in PDBx/mmCIF format. Benchmarking on the complete Protein Data Bank archive showed that MMseqs2 reduced archive-scale UniProtKB searches from hours with BLASTP to minutes, approximately 22-36 times faster, while curated mappings were recovered at top rank in 93.1% of cases. The remaining discrepancies mainly involved biologically ambiguous cases such as highly conserved proteins, chimeric constructs, or closely related orthologs. These results show that PDBe-SIFTS enables fast mapping, improving structural coherence in residue-level alignments while delivering the most up-to-date and accurate mappings, comparable to expert curation.

Tool: https://github.com/PDBeurope/SIFTS

Quick start notebook with example: https://github.com/PDBeurope/SIFTS/tree/master/notebooks

**Broader audience statement:** Matching protein sequences to their three-dimensional structures, and mapping annotations across both, is essential for understanding protein function, interactions, and molecular mechanisms. This integrated view enables richer interpretation of biological data and underpins advances in drug discovery, disease research, and protein engineering. PDBe-SIFTS provides an open and functional framework for structure–sequence mapping, allowing researchers and databases to run, inspect, and extend these mappings locally, while benefiting from faster searches, transparent scoring, and structurally informed residue-level alignments.

**Graphical abstract:** 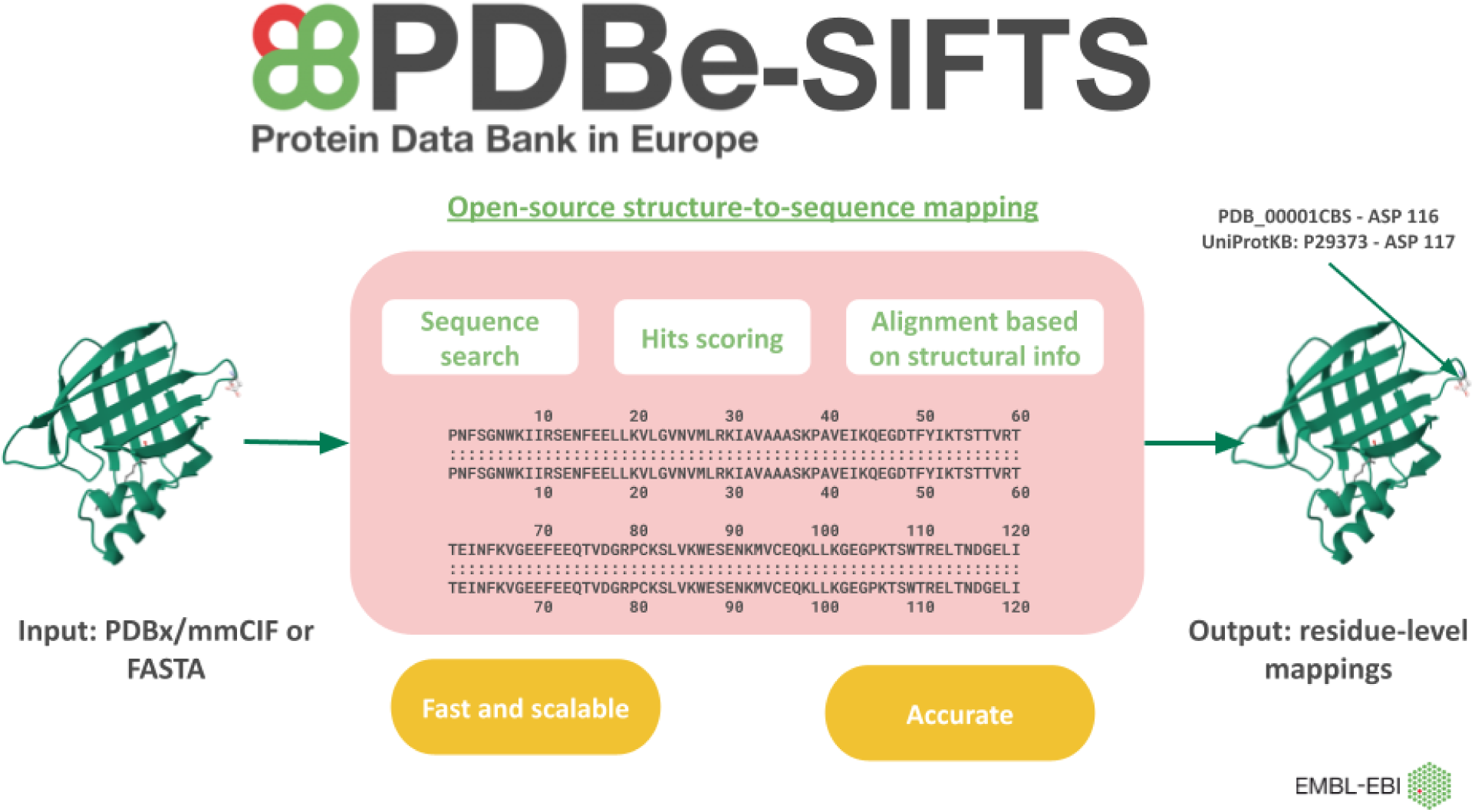

## Introduction

The rapid expansion of biological sequence and structural data has fundamentally transformed the landscape of structural bioinformatics, shifting the field from large-scale data generation and benchmarking towards structure-aware machine learning approaches that exploit 3D context at the residue level (Durairaj et al., 2023; Gutmanas et al., 2013; Samish et al., 2015; Tunyasuvunakool et al., 2021). UniProt KnowledgeBase (UniProtKB) continues to expand rapidly, growing from more than 227 million sequence records in release 2022_03 to approximately 246 million in release 2024_04 (The UniProt Consortium, 2023, 2025). The Protein Data Bank (PDB) has steadily increased in both size and structural complexity from 150 000 structures in 2019 to almost 250 000 in 2026 (Armstrong et al., 2020; wwPDB consortium, 2019). As a result, maintaining accurate and scalable integration between sequence and structure resources has become increasingly challenging.

Reliable mapping between UniProtKB sequences and PDB structures is a fundamental step in data integration and aligns with large-scale infrastructure initiatives such as ELIXIR (Harrow et al., 2021) and 3D-Bioinfo (Orengo et al., 2020). Such mappings enable the transfer of functional annotations, domain definitions, sequence variants, and disease-associated mutations onto three-dimensional structures (Martin, 2005). They also provide a common sequence-based residue numbering framework across multiple structural representations of the same protein including experimentally determined structures, predicted models, and simulations while enabling visualisation systems to accurately display these annotations (Choudhary et al., 2023). This, in turn, facilitates consistent interpretation and comparison across datasets.

The Structure Integration with Function, Taxonomy and Sequences (SIFTS) resource (Dana et al., 2019; Velankar et al., 2013) was developed in 2002 to address this need by providing residue-level mappings between UniProtKB and PDB entries. Over the years, SIFTS has become a central component of data integration within PDBe (Armstrong et al., 2020), UniProtKB (The UniProt Consortium, 2025), PDBe-KB (PDBe-KB consortium, 2020), and the PDB NextGen Archive (Choudhary et al., 2024).

The residue-level mapping from SIFTS is also incorporated into PDBx/mmCIF (Choudhary et al., 2023) through dedicated SIFTS categories in the PDBx/mmCIF dictionary (Westbrook et al., 2022). More broadly, it has served as a residue-mapping and annotation framework for a range of bioinformatics resources, including PDBsum (Laskowski et al., 2018), ProtVar (Stephenson et al., 2024), PDBj (Kinjo et al., 2018), RCSB PDB (Burley, Bhikadiya, et al., 2022), PLINDER (Durairaj et al., 2024), PDBRENUM (Faezov & Dunbrack, 2021), InterPro (Mitchell et al., 2019), Pfam (Finn et al., 2014), MobiDB (Piovesan et al., 2021), EIIpred (Liebold et al., 2026), AF2BIND (Gazizov et al., 2026) and tools designed to associate structural data with the scientific literature (Vollmar et al., 2026). However, the continued growth of both UniProtKB and the PDB archive poses ongoing challenges for scalability, computational efficiency, and handling of increasingly complex structural data. Because these underlying resources are dynamic, with new entries released and existing records revised or obsoleted over time, the mappings between them must be updated regularly. SIFTS data is regenerated weekly to ensure up-to-date cross-references and is accessible through the PDBe website, FTP, and API services.

SIFTS was developed as a collaborative resource between UniProtKB and PDBe to establish reliable and up-to-date residue-level mappings between UniProtKB sequences and PDB structures. In the early stages of the SIFTS project, co-location at EMBL-EBI helped kick-start development by enabling the PDBe and UniProtKB teams to leverage shared resources, internal databases, and tightly integrated infrastructure for rapid data exchange. As a result, the mapping workflow became tightly coupled to the internal infrastructure and was therefore not available as reusable research software outside the EMBL-EBI environment. In response to growing demand for local execution, these internal dependencies have since been removed, and a fully open-source, locally deployable implementation of this core SIFTS mapping framework is made available, enabling users to produce, inspect, and extend mappings for structures or sequences not available in the PDB and UniProtKB, respectively.

As research software becomes increasingly central to life-science data integration, transparent, accessible, interoperable, and reusable implementations are essential for verification, reuse, and long-term sustainability (Bayarri et al., 2024; Malik-Sheriff et al., 2020; Patel et al., 2023; Sansone et al., 2019; Wilkinson et al., 2016). For resources such as SIFTS, the need for local deployment, interoperability, and transparency is further underscored by the practical difficulty of reimplementing complex mapping frameworks from the published literature alone (Ehmki et al., 2026). Emerging initiatives such as the Molecular Dynamics Data Bank (MDDB) require robust SIFTS mappings to connect molecular dynamics simulations to UniProtKB and PDB annotations (Amaro et al., 2025).

Thus, we present the fully open-source, locally deployable implementation of the mapping framework underlying the SIFTS resource. In addition to making the pipeline openly available, the framework introduces several methodological improvements. Building on the previously described SIFTS scoring framework (Velankar et al., 2013), the ranking criteria is updated based on benchmarking manually curated mappings. It aims to prioritise biologically meaningful mappings while maintaining consistency across related entries. A scalable sequence-search strategy based on MMseqs2 (Steinegger & Söding, 2017) is implemented, enabling efficient processing of large datasets. Finally, the residue-level mapping procedure was refined by incorporating structural connectivity constraints, ensuring that the resulting mapping reflects experimentally observed backbone connectivity and reduces alignment artefacts.

Tool: https://github.com/PDBeurope/SIFTS

Quick start notebook with example: https://github.com/PDBeurope/SIFTS/tree/master/notebooks

## Results

### Pipeline and software design

The entire codebase is now available in the open-source PDBe-SIFTS repository (https://github.com/PDBeurope/SIFTS/). PDBe-SIFTS is distributed as a standard, pip-installable Python package (v1.0, Python ≥ 3.10). A command-line interface (CLI) exposes six subcommands: build_db, fasta_build, sequence_match, segments, db_load, and sifts2mmcif, covering the full package from database construction to generating residue-level mapping in PDBx/mmCIF (Westbrook et al., 2022), including SIFTS annotations (Choudhary et al., 2023). Each module can be executed independently and has its own input requirements. These subcommands are illustrated in Figure 1 and are detailed below:

**Figure 1.**
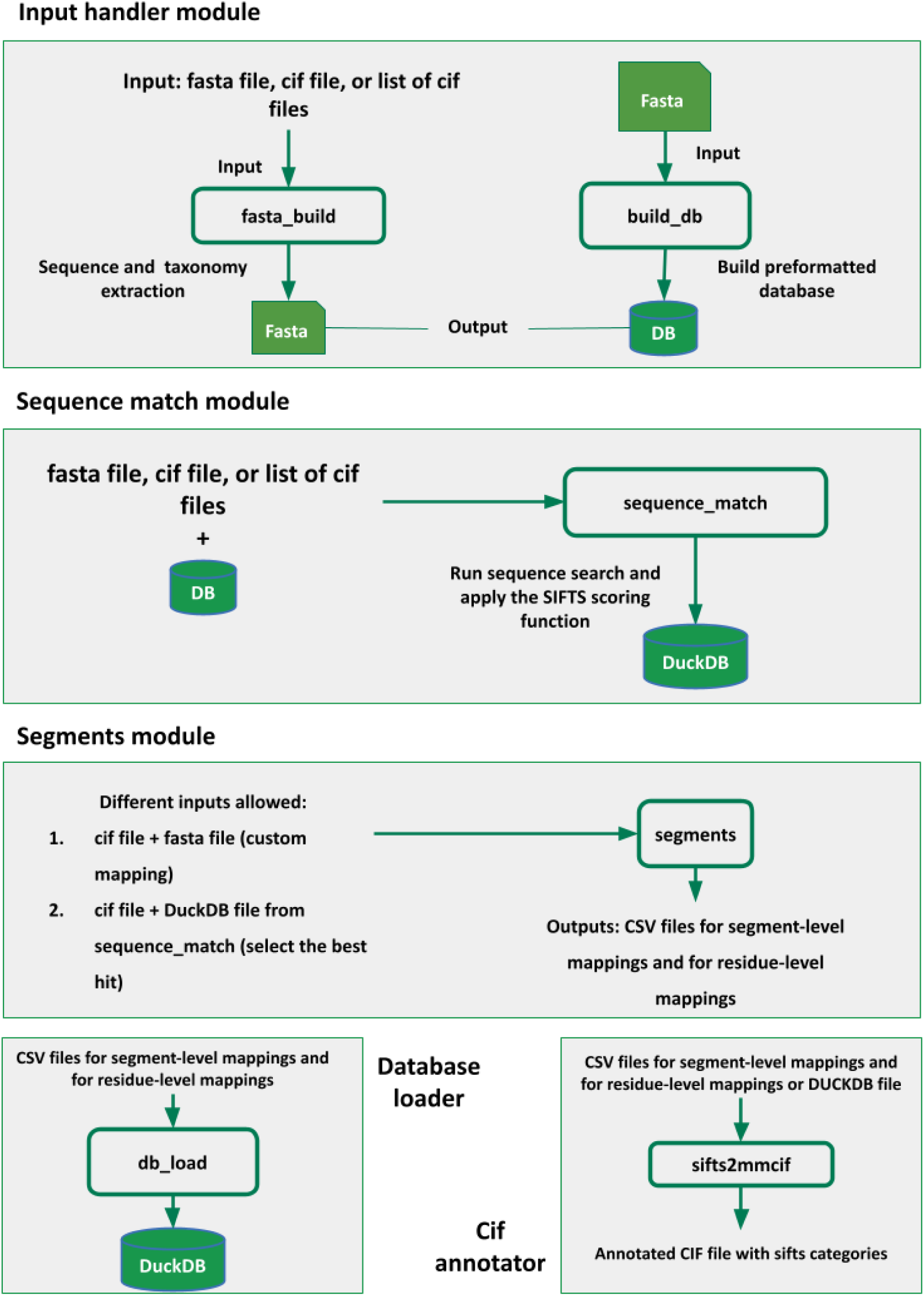
Overview of the main pipeline of PDBe-SIFTS. Modules fasta_build, build_db, sequence_match, segments, db_load, and sifts2mmcif are used through CLI commands.

#### (1) build_db

Constructs an indexed reference sequence database from a user-supplied FASTA file following UniProtKB FASTA conventions (https://www.uniprot.org/help/fasta-headers). Two software suites are supported for sequence searching: MMseqs2 (Steinegger & Söding, 2017) and BLASTP (Camacho et al., 2009). At this stage, a taxonomy-mapping file, provided as a tab-separated list of sequence identifiers and NCBI taxon identifiers, is required to enable taxonomy-aware scoring in downstream steps.

#### (2) fasta_build

Parses one or more mmCIF files to extract amino acid sequences for each protein structure chain and writes them to a multi-FASTA file. Sequences are derived from the _entity_poly category, which stores the sample polymer sequence independently of coordinate completeness. This approach ensures that unobserved residues in the structure, such as flexible loops, are properly represented, avoiding incorrect alignments that can occur with standard sequence-alignment methods. The resulting FASTA file can be used as a custom query set for the sequence_match step.

#### (3) sequence_match

Compares the sequences from the fasta_build step or sequences provided as a FASTA file against the reference database generated by **build_db** and identifies the highest-scoring match for each polypeptide chain. The scoring scheme applies a 90% sequence-identity cutoff to account for sequence differences, in combination with the other metrics described in the Methods section. Ranked hits and residue-level mappings are stored in a DuckDB database to support scalable queries over large datasets. DuckDB provides an embedded, columnar analytics engine with a single-file database format and efficient vectorised execution, enabling hundreds of millions of rows of mapping data to be queried and aggregated with modest memory requirements and without a separate database server. This design enables users to run archive-scale evaluations and downstream analyses of SIFTS mappings directly from Python.

#### (4) segments

This module performs residue-level alignment and segment generation (see Methods) for each chain in the PDBx/mmCIF file and the identified reference sequence from the sequence_match step. The module also provides an option to bypass the database search step and use the user-supplied sequence as a reference sequence directly for segment generation and residue-level mapping.

The protein sequence for the structure is extracted from _pdbx_poly_seq_scheme (if missing this is automatically added by the module using sample sequence in _entity_poly or a user-provided input) is aligned to the reference sequence using Lalign (Huang & Miller, 1991), which implements the Smith–Waterman local alignment algorithm and reports multiple non-overlapping local alignments. To ensure accurate mapping, each chain is partitioned into segments identified as continuous stretches of residues that share the same taxonomic identifier. Each segment mapped to a reference sequence is then used to produce residue level mappings. The outputs are written as gzip-compressed CSV files for segment-level mappings and for residue-level mappings, both stored within an entry-specific output.

#### (5) db_load

Performs bulk import of the segment and residue mapping files generated by the **segments command** into the DuckDB tables sifts_xref_segment and sifts_xref_residue. It scans a root output directory, identifies valid entry-level files, and appends their contents in a single transactional operation, thereby enabling efficient cross-entry queries following large-scale processing. The DuckDB file contains three tables, and the generated CSV files follow the same column logic:

1. **Hits (only in DuckDB).** This table contains candidate sequence hits produced during the sequence match step and stored only in DuckDB. Each row corresponds to one structure entity pair matched against a reference sequence, together with the alignment and scoring information used to rank the hit. It contains the following columns: entry (structure id), entity (chain id), accession (reference sequence id), alignment_len, query_len, mismatch, query_start, query_end, target_start, target_end, e-value, bit_score, identity, coverage, query_aligned, target_aligned, target_tax_id, query_tax_id, sifts_score, pdb_cross_references, adjusted_score, tax_score, dataset_score, hit_rank.
2. **sifts_xref_segment**. This table stores the segment-level mapping output, with one row per aligned segment. It describes the continuous aligned blocks used to map residues, including entry, entity, auth_asym_id, struct_asym_id, accession (reference sequence id), name, seq_version, query_start, query_end, target_start, target_end, auth_start, auth_end, auth_code, target_alignment, query_alignment, identity, score, conflicts, modifications, best_mapping (bool), cannonical_acc (bool) and chimera (bool).
3. **sifts_xref_residue.** This table contains the residue-level mapping output, with one row per residue. It provides the final one-to-one residue correspondence between the two sequences. It includes: entry, entity, auth_asym_id, struct_asym_id, target_segment_id, auth_seq_id, auth_seq_id_ins_code, query_seq_id, target_seq_id, observed, query_one_letter_code, target_one_letter_code, chem_comp_id, type, tax_id, best_mapping (bool), canonical_acc (bool), residue_id.

#### (6) sifts2mmcif

Integrates the computed SIFTS annotations into the source PDBx/mmCIF file by populating the “_pdbx_sifts_unp_segments” and “_pdbx_sifts_xref_db” categories (Choudhary et al., 2023). Additionally, the “atom_site” category is extended to include the sequence accession and residue number from the best available mapping, enabling each atomic coordinate to be associated directly with the corresponding mapped residue. These files are also compatible with visualisation software such as Mol*, allowing users to visualise both reference sequence numbering and structure residue numbering simultaneously, as shown in Figure 2 where the residue ASP 116 (PDB 1CBS, Kleywegt et al., 1994) is mapped to the ASP 117 of the UniProt accession P29373.

**Figure 2.**
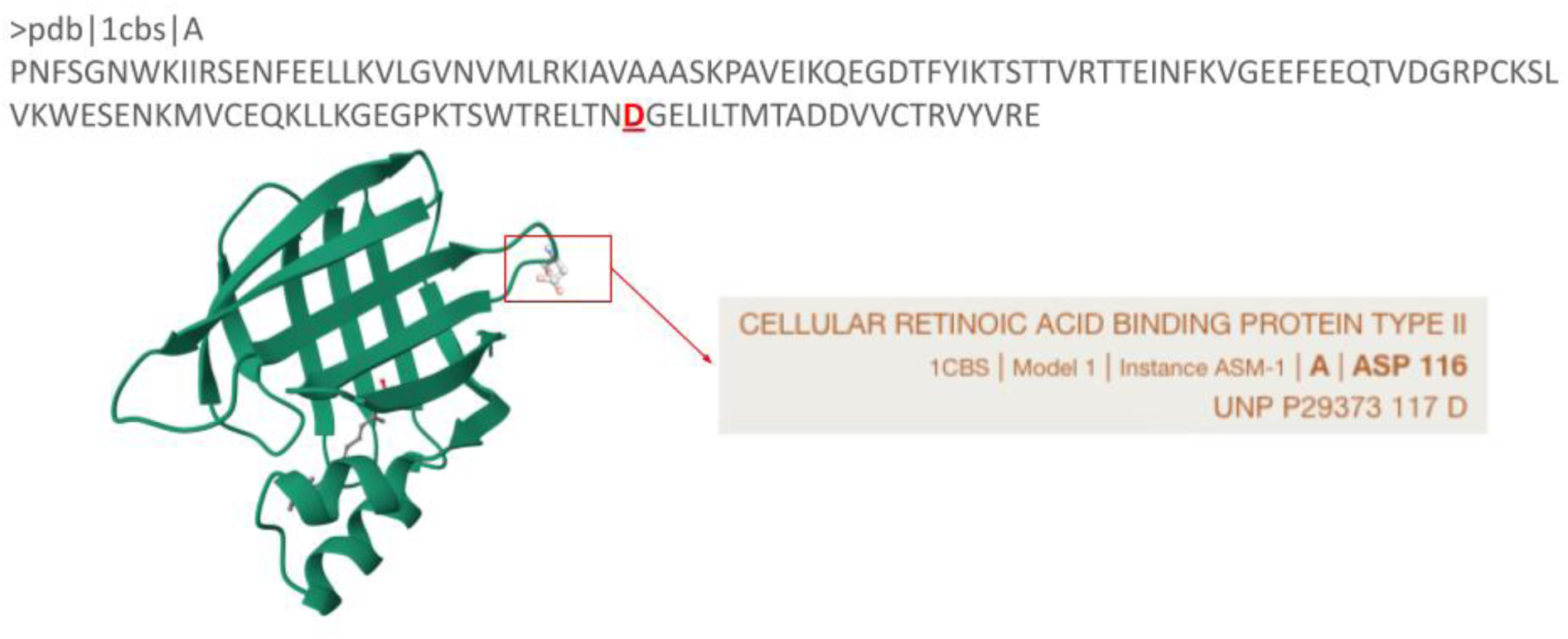
Mol* visualisation of PDB structure 1CBS using the PDBx/mmCIF file with integrated SIFTS annotations showing the correspondence between ASP 116 in the PDB structure and ASP 117 in UniProt accession P29373.

### Benchmarking UniProtKB-PDB mappings

#### Runtime Performance of BLASTP and MMseqs2

We investigated and benchmarked two input modes: (i) multiple PDBx/mmCIF structures as input, and (ii) a single PDBx/mmCIF structure as input. These two settings were evaluated separately to assess performance and scalability under single- and multi-queries conditions, as these modes impose different computational characteristics and optimisation requirements.

#### Multiple queries

The performance of the sequence_match (UniProtKB-PDB sequence mapping) component was evaluated along two axes: speed and accuracy. The benchmark was performed on the complete PDB archive (see Methods), using either UniProtKB/SwissProt (curated dataset: ∼575,000 sequences) or the full UniProtKB (curated + non-reviewed entries: ∼ 200 million sequences) sequence database, with all PDB structures as input. Table 1 summarises the search time for BLASTP and MMseqs2 (easy-search) when querying PDB sequences against UniProtKB/Swiss-Prot or UniProtKB. MMseq2 was evaluated at two different sensitivity levels (5.7 and 7.5) to assess the trade-off between search thoroughness and runtime. The sensitivity parameter controls search depth: higher values enable detection of more distant or ambiguous matches but at the cost of increased runtime, while lower values favour faster execution with a slight reduction in sensitivity. The default setting of 5.7 provides a balanced compromise between performance and sensitivity.

**Table 1.** MMseqs2 and BLASTP search times for the entire PDB archive against UniProtKB/Swiss-Prot and UniProtKB. UniProtKB contains over 200 million sequences, whereas the PDB contains nearly 530 000 sequences. The hit ranking columns represent the time to apply the scoring function to all matches, independent of the tool used.

| Database | MMSeqs2 | BLASTP | Hit ranking |
| --- | --- | --- | --- |
| UniprotKB/Swiss-Prot | ~ 10 minutes (sensitivity: 5.7)<br>~ 16 minutes (sensitivity: 7.5) | ~ 6 hours | ~ 4 minutes |
| UniProtKB | ~ 6 hours (sensitivity: 5.7)<br>~ 34 hours (sensitivity: 7.5) | N/A | ~ 30 minutes |

MMseqs2 processes all Swiss-Prot searches in approximately 10 minutes (sensitivity 5.7) or 16 minutes (sensitivity 7.5), compared to roughly 6 hours for BLASTP. For UniProtKB searches across the whole PDB archive, MMseqs2 requires approximately 6 hours (sensitivity 5.7) or 34 hours (sensitivity 7.5). It should be noted that in the context of SIFTS, where the focus is on identifying exact or closest matches rather than distant homologs, a lower sensitivity setting of 5.7 is considered sufficient and appropriate. Equivalent BLASTP archive-scale searches against UniProtKB were not performed due to computational constraints.

#### Single queries

To evaluate performance in the single-query setting, benchmarking was performed separately using individual PDBx/mmCIF structures as input, in contrast to the multi-structure benchmarking described above, where all PDB structures were given as input. MMseqs2 is primarily optimised for many-to-many comparisons, whereas efficient single-to-many searches require minimising computational overhead. To optimise single query searches, an optional gapless prefilter (as implemented in MMseqs2) was introduced to accelerate searches against large databases by restricting the initial stage to ungapped k-mer matches, thereby reducing the number of candidate alignments. Thus, the sequence_match was benchmarked on 150 PDB entities spanning a wide range of sequence lengths, from 5 to 8,572 residues (See Materials and methods - Single queries).

Search time linearly increases with sequence length (Figure 3). For short sequences (50–100 residues), the mean search time was approximately 39 seconds, reflecting a near-constant overhead dominated by database initialisation. For sequences in the 1,500–2,000 residue range, mean search time rose to ∼142 seconds, and reached ∼347 seconds for the 5,000–6,000 residue bin. The single entry exceeding 8,000 residues (PDB entry: 9b4h, chain X, 8,572 residues) completed in 767 seconds. Standard deviation within bins is generally low but increases in long sequence bins. Overall, the search time remains low with a median below 6 minutes across all benchmarked entities (13 minutes for the longer one).

**Figure 3.**
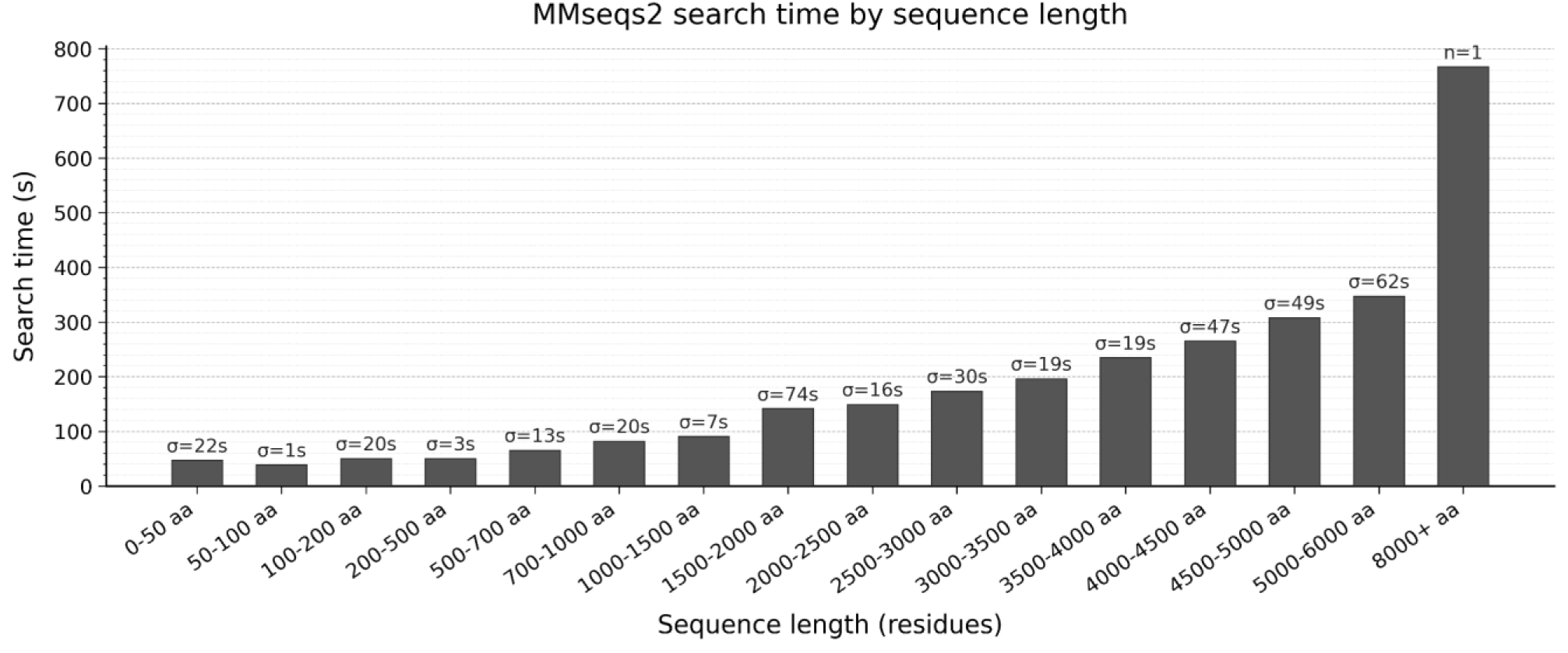
Computational performance of sequence_match across 150 PDB entries, grouped by sequence length. Mean search time per bin is shown in seconds, and standard deviation is indicated for bins with more than one entry.

#### Accuracy of sequence matches

The original SIFTS scoring scheme (Velankar et al., 2013) was refined to align with the latest UniProtKB and PDB data standards and to reflect up-to-date biocuration practices by PDB (wwPDB consortium, 2019) and UniProtKB (The UniProt Consortium, 2025) annotators. A heuristic and interpretable composite score was developed (details in Methods) that integrates sequence similarity, mismatch penalisation, taxonomy matching and dataset provenance to prioritise matches that closely approximate expert curation while minimising the need for extensive manual intervention wherever possible.

To evaluate scoring scheme performance, the ranking of manually curated mappings (SIFTS data from January 2026) was compared to the automatically identified match for each PDB chain in the curated dataset (See Methods). As shown in Figure 4, the curated mapping matched the top ranked hit in 93.1% of cases, while 6.9% yielded an alternative top-scoring mapping. In the latter cases, the curated mapping was typically ranked between 2 and 5. The remaining cases largely correspond to biologically ambiguous situations, including highly conserved proteins (such as ribosomal proteins), engineered chimeric constructs (also referred to as fusion proteins, which are made by combining parts from two or more distinct proteins into a single new, hybrid protein, or proteins with very similar orthologs across species. In such scenarios, multiple UniProtKB accessions may represent equally plausible mappings from a purely sequence-based perspective. Notably, even in these cases, the process consistently ranks the curated mappings in the top 10 matches.

**Figure 4.**
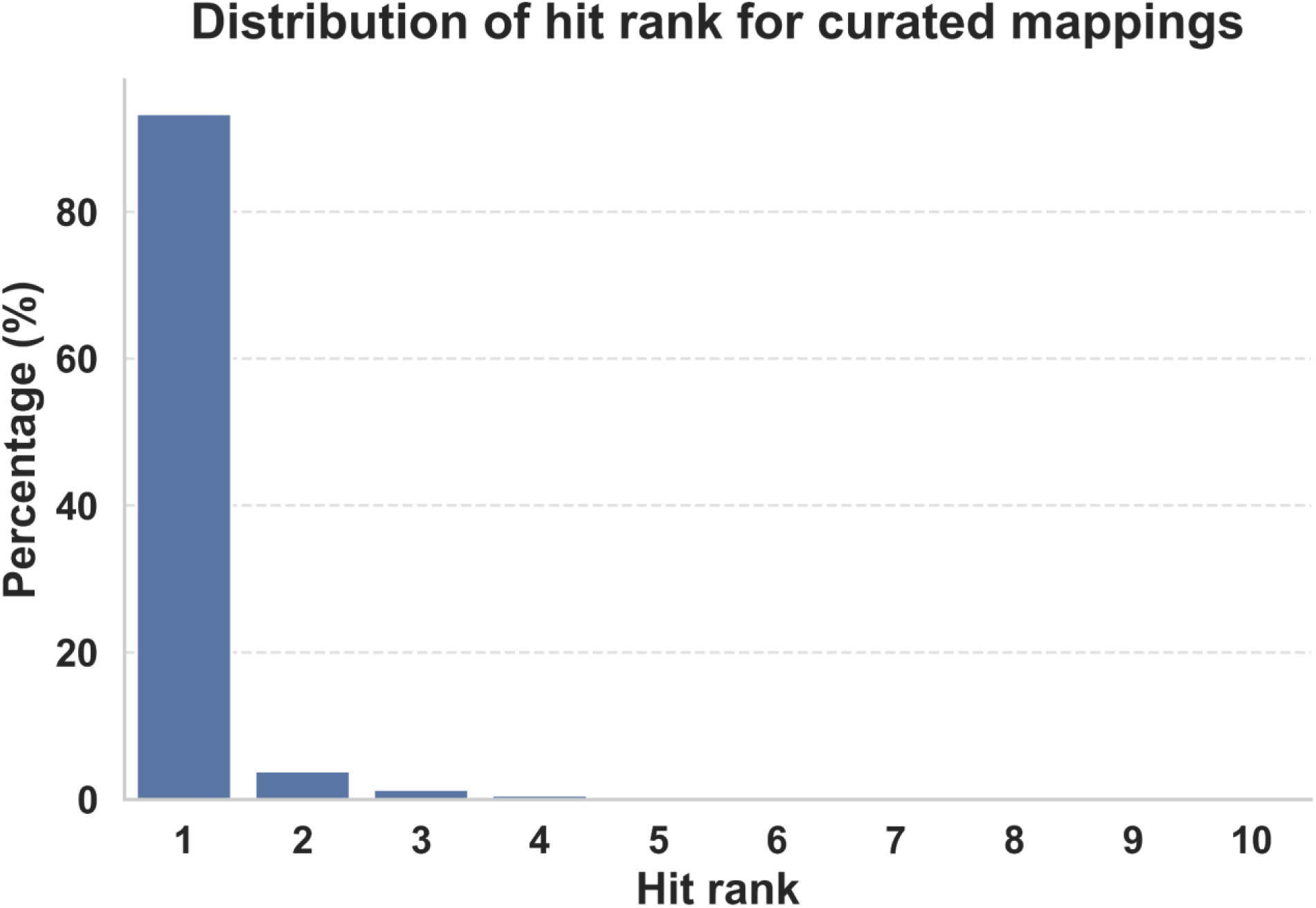
Recovery of curated UniProtKB-PDB mappings at different ranks.

### Refinement of sequence-structure alignments using structural connectivity

Sequence-structure mappings are commonly generated using classical pairwise sequence alignment algorithms. Although generally effective, these methods can introduce artefacts when experimentally determined structures contain unresolved regions or regions of low sequence complexity or ambiguity, where multiple alignments yield similar scores. In latter cases, alignment algorithms, depending on the input parameters, may optimise the score by inserting short, isolated matches within long gaps, leading to residue correspondences that are inconsistent with the actual structural connectivity of the protein backbone.

An illustrative example is shown in Figure 5, which shows the PDB structure 2XDE (Blair et al., 2010), chain A. In this structure, residues 87–96 are not observed in the experimentally modelled coordinates relative to the UniProtKB sequence P12497, reflecting a deletion.

**Figure 5.**
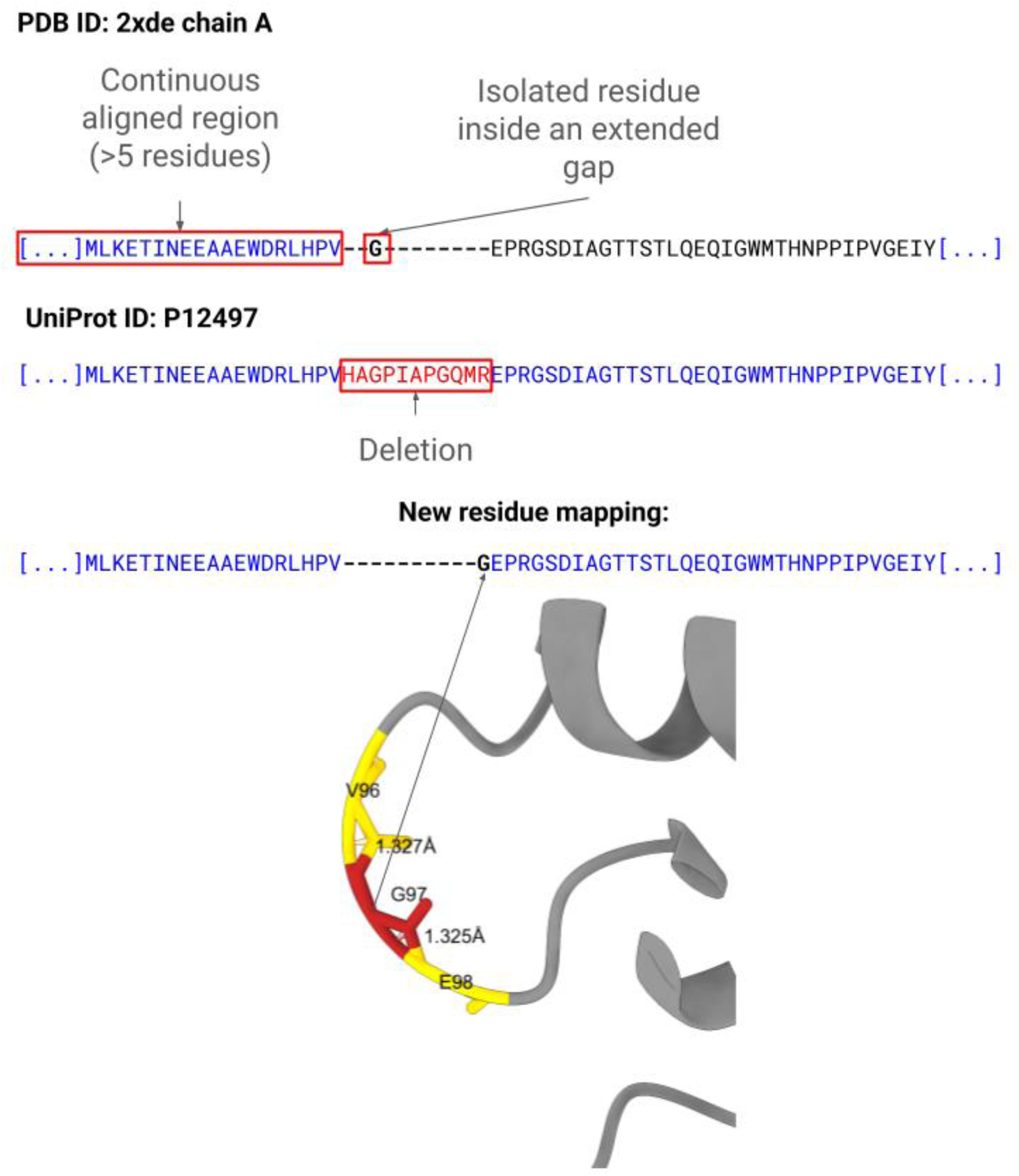
Backbone connectivity refines SIFTS residue-level alignments by merging isolated residues within extended gaps into adjacent continuous aligned regions. Blue indicates aligned regions, red indicates deletions in the PDB sequence relative to UniProtKB, and the example is derived from PDB structure 2XDE.

In the original sequence–structure alignment, a single glycine residue is incorrectly aligned within this missing segment. This results in an isolated match surrounded by gaps, artificially fragmenting what should be represented as a single continuous unmapped region. Consequently, the alignment no longer reflects the true structural connectivity of the protein.

The connectivity-based refinement addresses this artefact by enforcing consistency with backbone connectivity derived from the three-dimensional structure. The refined alignment removes the isolated match and represents the unresolved region as a single contiguous gap.

This approach restores a structurally consistent mapping between sequence and experimentally observed structure. At the scale of the PDB archive, approximately 2% of chain alignments are affected by this issue (defined as the proportion of impacted chains among all chains).

## Discussion

In this work, we present an updated framework for structure-to-sequence mapping that combines scalable sequence search with an interpretable scoring scheme and a structurally informed residue-level refinement strategy to derive final mapping. Together, these improvements enhance mapping accuracy, computational efficiency, and structural consistency, enabling more robust SIFTS annotations across the entire PDB archive. These developments are particularly important given the continuous growth in the size and complexity of the PDB and UniProtKB databases.

PDBe-SIFTS is now a fully open-source software package, allowing users to run the SIFTS pipeline locally on custom sequence or structure datasets. Making the mapping workflow available as a standalone research software improves transparency and reuse across independent computational environments (Martín del Pico et al., 2024) while enabling large-scale sequence-structure integration across independently generated datasets.

The benchmarking results demonstrate that the updated SIFTS pipeline can process archive-scale sequence searches efficiently using MMseqs2 while maintaining high mapping accuracy. Curated UniProtKB PDB mappings were recovered as top-ranked hits in much of the archive, confirming that the scoring scheme effectively prioritises best matches. The bounded and interpretable scoring formulation also makes mapping decisions more traceable in large-scale automated analyses.

Despite these improvements, accurate automatic mapping remains challenging for chimeric or engineered fusion proteins (Baldo, 2015) in the absence of manual curation. Although such entries currently represent only a small fraction of the PDB archive, their prevalence is expected to increase with the growing use of engineered constructs in structural biology and therapeutic design (Marsh & Owen, 2023; Silver et al., 2021). Similarly, cases involving highly conserved proteins, or closely related orthologs, may lead to ambiguous structure-sequence mappings requiring additional biological context to determine the most appropriate match.

Future developments may therefore focus on integrating additional sources of information, such as experimental annotations or construct metadata, to further improve the automated identification of complex protein architectures. The modular design of PDBe-SIFTS could support future extensions to important structural contexts such as antibody or peptide complexes and nucleic acid-containing assemblies, which are both well represented in the structural biology ecosystem (Burley, Berman, et al., 2022; Dunbar et al., 2014; Lee et al., 2022).

Overall, PDBe-SIFTS provides a scalable and reproducible framework for sequence–structure integration, enabling reliable residue-level mappings between UniProtKB and the PDB while supporting the growing needs of modern structural bioinformatics infrastructures.

## Materials and methods

From the previously described SIFTS procedures (Dana et al., 2019; Velankar et al., 2013), the following two main components have been modified in the current implementation.

### 1. Sequence match

The first component (sequence_match) is the automated process used to identify structure to sequence matches. The updated implementation introduces two main improvements **(1)** the ability to perform scalable and fast sequence searches using MMseqs2 (Steinegger & Söding, 2017) or the use of BLAST (Camacho et al., 2009); and **(2)** a revised scoring scheme for ranking hits, defined in Figure 6. PDBe-SIFTS ranking scheme is a heuristic and interpretable composite score that combines sequence similarity, mismatch penalisation, taxonomy/lineage match, and dataset provenance (for UniProtKB) to prioritise biologically meaningful matches.

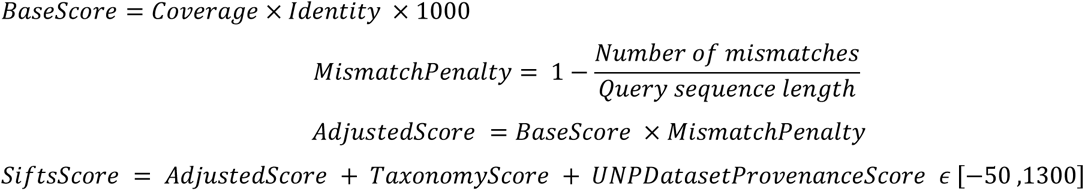

Hits are ordered by SiftsScore and number of PDB cross-references

**Figure 6.**
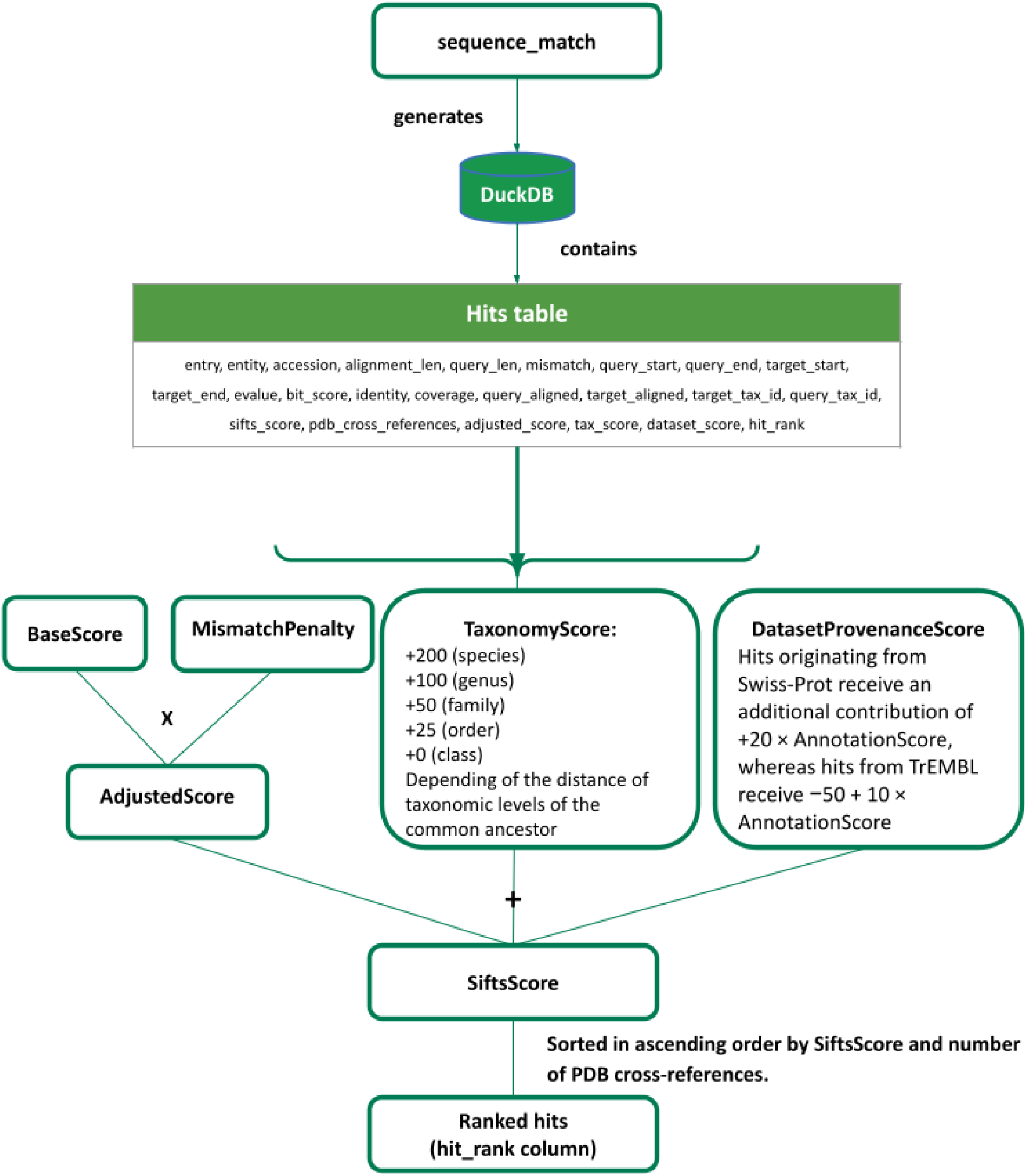
SIFTS scoring scheme and ranking process.

The BaseScore is calculated from coverage (the fraction of the protein structure sequence aligned to the reference sequence) and Identity (the fraction of aligned residues that are identical between the protein structure and the reference sequence). The BaseScore is multiplied by 1000 to scale it to a numerical range comparable to the additional weighting terms while ensuring that sequence similarity remains the dominant contributor to the final score (SiftsScore).

To account for sequence divergence, the number of mismatches (positions in the alignment where residues differ, excluding gaps) is used to compute a MismatchPenalty. This penalty ensures that a few mismatches in a short sequence strongly reduce the score, whereas the same number of mismatches in a long sequence has a proportionally smaller effect, providing a fair measure of sequence divergence across different alignment lengths. The AdjustedScore, therefore, represents a refined estimate of sequence similarity after accounting for mismatches.

The FinalScore is obtained by combining the AdjustedScore with additional biologically motivated criteria. Taxonomic score represents the most important of these criteria. Additional +200 points are assigned when taxonomies of the query and the target exactly match at the species level. Reduced contributions (+100, +50, +25) are assigned for matches at the genus, family and order levels, corresponding to one, two, or three taxonomic levels of the most recent common ancestor, respectively. This taxonomic distance is computed using the tool ETE 3 (Huerta-Cepas et al., 2016).

Dataset provenance represents the second additional criterion. Hits originating from UniProtKB/Swiss-Prot receive an additional contribution of (+20 × AnnotationScore), whereas hits from UniProtKB/TrEMBL receive (−50 + (10 × AnnotationScore)). The AnnotationScore corresponds to the UniProtKB annotation score, which provides a heuristic estimate of the level of annotation available for each entry. The scoring scheme can, however, be applied to non-UniProtKB datasets by removing these scores.

The ranking protocol was designed to reflect the main criteria typically used during manual curation of UniProtKB-PDB mappings while remaining simple and interpretable. Taxonomic similarity represents the second most important criterion. Exact species matches receive the highest additional weight, while progressively smaller contributions are assigned to matches at higher taxonomic levels. This reflects the biological expectation that protein structures should preferentially map to sequences originating from the same organism whenever possible. Finally, dataset provenance is incorporated to favour UniProtKB entries with higher annotation reliability. UniProtKB/Swiss-Prot (curated) entries receive a higher contribution due to their manual curation, whereas UniProtKB/TrEMBL (not curated) entries receive a smaller weight proportional to their UniProtKB annotation score.

It is common for multiple candidate hits to receive identical scores. In such cases, the UniProtKB accession with the most PDB cross-references is selected. This tie-breaking strategy ensures that protein structures for the same protein are consistently mapped to the same accession. The number of PDB cross-references is intentionally excluded from the scoring function to keep the score bounded and comparable across proteins with varying levels of structural representation in the archive.

The resulting original and ranked hits are saved as TSV files and in a DuckDB (Raasveldt & Muehleisen, 2018/2026) database. These files serve as the starting point for generating residue-level correspondences between the protein structure and the selected hit sequence for each mapped protein chain.

### 2. Residue-level mapping

The second component (segments) is an automated process that generates residue-level mappings between the reference sequence and the corresponding protein structure entity. A molecular entity is a chemically distinct component of the structure as represented in the PDBx/mmCIF file and is of three types: polymer, non-polymer or water. This component takes as input the highest-scoring hit (including the one with the highest number of PDB cross-references) and performs a local alignment that accounts for structural features such as linkers, expression tags, mutations, deletions, and additions. It can also take a custom hit from a FASTA file as input. This alignment is computed using *lalign36* (Huang & Miller, 1991). From this, the SIFTS segments are generated. These segments represent continuous, connected regions of the structure’s chain with the same taxonomy identifier or accession and serve as the fundamental building blocks of the residue-level mapping. According to this definition, several checks are performed during the generation of SIFTS segments to ensure structural and biological consistency:

- all residues within a segment share the same taxonomy identifier or accession;
- residues are connected at the backbone level (connectivity check).

Sequence alignments may introduce artificial gaps that do not correspond to real structural breaks. To address this issue, structural backbone connectivity is used to refine residue-level mappings and preserve the continuity of experimentally observed protein chains. Residues are classified into two categories (see Figure 5): (1) residues belonging to continuous regions of more than five residues; and (2) residues located in extended gaps, defined as regions where more than one consecutive gap symbol separates aligned residues.

In extended gaps, isolated residues (<5) may occasionally appear. When a residue remains physically connected to the neighbouring structural residue, the gap pattern reflects an artefact of the sequence alignment rather than a true structural discontinuity. In these cases, the residue is merged into the adjacent segment to maintain structural coherence. The connectivity assessment between two residues relies on the distance between the backbone C atom of the preceding residue and the backbone N atom of the following residue. Two residues are considered connected when this distance is below a threshold of 1.42 Å. This value allows for small coordinate deviations observed in experimental structures while remaining consistent with the typical peptide bond length (∼1.33 Å), thereby providing a conservative tolerance that preserves genuine backbone connectivity and avoids artificial discontinuities. The identification of the relevant backbone atoms is guided by the Chemical Component Dictionary (CCD) (Westbrook et al., 2015) files available through the PDBe FTP repository (ftp.ebi.ac.uk/pub/databases/msd/pdbechem_v2/ccd/components.cif). This approach allows SIFTS to determine the appropriate N-terminal and C-terminal atoms for each residue directly from the CCD definitions when evaluating structural connectivity.

### UniProtKB-PDB mappings

#### Multiple queries

To evaluate both the speed and accuracy of the first component (sequence match), we used the complete set of SIFTS curated mappings available from the EMBL-EBI FTP repository (January 2026; https://ftp.ebi.ac.uk/pub/databases/msd/sifts/ or in the repository https://github.com/PDBeurope/SIFTS/blob/master/tests/data/curated_mappings/curated.csv]). These curated mappings link PDB entry–chain pairs to UniProtKB accessions. The dataset comprises approximately 380K curated mappings and spans a wide range of mapping scenarios, including chimeric proteins, highly conserved sequences, UniProtKB/Swiss-Prot or UniProtKB mappings, large macromolecular assemblies such as ribosomes, sequences of unknown origin, UniProtKB reference proteome entries, mappings with taxonomic mismatches, and cases with both low and high sequence coverage. UniProtKB/TrEMBL (unreviewed) contains protein sequences with automatically generated annotations from large-scale computational analyses, whereas UniProtKB/Swiss-Prot (reviewed) is a high-quality, non-redundant database of manually curated protein sequences integrating experimental evidence, computed features, and expert interpretation.

#### Single queries

To evaluate the computational performance of the pdbe_sifts sequence_match, a set of 150 PDB protein entities was selected to provide uniform coverage across a range of sequence lengths, from fewer than 50 to more than 8,000 residues (list below respecting the format PDBID_ENTITY). Entities were assigned to bins of increasing sequence length, with ten entries per bin where possible, ensuring that the benchmark captures performance across the full diversity of chain sizes encountered in the PDB. All mappings were performed against UniProtKB [January 2026]. Mean search time and standard deviation were computed per bin. No parallelism was used across entities; each run was executed sequentially as a single process. The list of the PDB and entity IDs is available in the repository: https://github.com/PDBeurope/SIFTS/tree/master/tests/data/curated_mappings/single_query.csv.

### Hardware and software specifications

All runs were performed using the EMBL-EBI CPU cluster on 1 node of 48 cores and 400 GB of memory. The fasta files “uniprot_trembl.fasta.gz” and “uniprot_sprot.fasta.gz”, downloadable from UniProtKB FTP (January 2026 released) were used to pre-build the database for each alignment tool.

### MMSeqs2 and BLASTP configurations

#### MMSeqs2

databases were built using the MMseqs2 modules createdb, createtaxdb, and createindex for UniProtKB Swiss-Prot and UniProtKB, respectively, resulting in two sets of files of 4 GB and 862 GB. The search was performed using the easy-search module (sensitivity 5.7 or 7.5, minimum sequence identity 0.9, all other parameters set to default).

#### BLASTP

databases were built using the makeblastdb module for UniProtKB Swiss-Prot and UniProtKB. Then the search was performed using blastp with default parameters.

## Code availability and dependencies

The PDBe-SIFTS pipeline is available as an open-source software at: https://github.com/PDBeurope/SIFTS. The package can be installed via PyPI or micromamba and executed locally.

PDBe-SIFTS depends on the following Python packages: numpy (Harris et al., 2020), gemmi (Wojdyr, 2022) (mmCIF parsing), duckdb (storage and data access), biopython (Cock et al., 2009) (alignment I/O), pymmseqs (MMseqs2 Python wrapper), requests and xml (UniProtKB API), pandas (McKinney, 2010) (data manipulation), scikit-learn (Pedregosa et al., 2011), omegaconf (configuration), ete3 (taxonomy), platformdirs, python-dateutil, pyyaml, funcy and coloredlogs (logging). The following external binaries must be installed separately and available on PATH: MMseqs2 (sequence search), lalign36 (FASTA36 package, local alignment), and BLASTP (NCBI BLAST).

## Acknowledgments

The authors thank the EMBL-EBI and the PDBe team for their support and contributions to this work. The authors also thank Milot Mirdita for assistance in optimising MMseqs2 parameters and in addressing technical challenges. This work was supported by the European Union’s Horizon Europe programme under grant agreement No. 101094651 of the MDDB Project. Conny Yu is supported by funding from the National Institutes of Health [U24HG007822 to UniProt].

## Data availability and accessibility statement

PDBe-SIFTS is an open-source computational tool for protein sequence-structure mapping. The source code is freely available at https://github.com/PDBeurope/SIFTS, and a quick-start notebook illustrating its use is available at https://github.com/PDBeurope/SIFTS/tree/master/notebooks. Public source data used in this study were obtained from the Protein Data Bank in PDBx/mmCIF format, SIFTS flatfiles and from UniProtKB releases described in the manuscript. The repository contains the benchmarked mappings here: https://github.com/PDBeurope/SIFTS/tree/master/tests/data/curated_mappings/curated.csv and https://github.com/PDBeurope/SIFTS/tree/master/tests/data/curated_mappings/single_query.csv.

## Affiliations and CRediT

- Adam Bellaiche (1): Conceptualitization, Data Curation, Validation, Project Administration, Visualization, Formal Analysis, Investigation, Methodology, Software, Writing – Original Draft Preparation
- Preeti Choudhary (1): Writing – Review & Editing, Conceptualization, Methodology, Software, Validation, Project Administration, Supervision
- Jennifer Fleming(1): Writing – Review & Editing, Project Administration, Supervision
- Sreenath Nair (1): Writing – Review & Editing, Software, Validation, Resources, Data Curation
- Sameer Velankar (1): Writing – Review & Editing, Conceptualization, Funding Acquisition, Project Administration, Supervision
- Conny Yu (2): Writing – Review & Editing, Validation, Resources, Investigation, Data Curation
- Syed Ahsan Tanweer (1) - Software
- Genevieve Laura Evans (1) - Investigation, Data Curation
- Stephanie W Lo (2) - Writing – Review & Editing
- Maria Martin (3) - Review & Editing

## Abbreviations and symbols

BLASTP: Basic Local Alignment Search Tool for Proteins
MDDB: Molecular Dynamics Data Bank
MMSeqs2: Many-against-Many sequence searching 2
PDB: Protein Data Bank
PDBe: Protein Data Bank in Europe
SIFTS: Structure Integration with Function, Taxonomy and Sequences
UniProtKB: UniProt KnowledgeBase

